# Functional traits and soil water availability shape competitive interactions in a diploid-polyploid complex

**DOI:** 10.1101/2024.08.23.608238

**Authors:** Alba Rodríguez-Parra, Javier López-Jurado, Enrique Mateos-Naranjo, Francisco Balao

## Abstract

1. Even within those polyploid plant species that become established initially, only a few persist in the long term. Competitive interactions between polyploids and their ancestral cytotypes in secondary contact zones can contribute to local extinctions. Environmental factors such as water availability and functional trait divergence may influence these interactions.
2. We conducted a greenhouse competition experiment with four cytotypes (2x, 4x, 6x, and 12x) of *Dianthus broteri* under two contrasting irrigation regimes. We estimated niche and fitness differences and predicted the pairwise competitive outcomes. Additionally, we explored the influence of leaf physiological functional traits (SLA, A_N_, g_s_, LDMC, F_v_/F_m_, and _i_WUE) on competitive interactions.
3. Soil water availability modified the competitive dynamics between cytotypes and predicted competitive exclusion. Under high water availability, lower ploidy levels (2x and 4x) outcompeted higher ploidy cytotypes (12x, 6x), while the latter exhibited greater competitive abilities under low water availability. These differences were explained by functional traits related to competitive effects (SLA) and competition tolerance (A_N_, F_v_/F_m_, and _i_WUE).
4. *Synthesis:* Our study emphasizes that the long-term fate of polyploids largely depends on water availability, with high polyploids having a competitive advantage in arid environments. This ultimately highlights the role of functional traits in shaping the competitive dynamics between cytotypes.

## Introduction

Polyploidy is a crucial mechanism in plant evolution (Grant, 1981; Soltis et al., 2009). Despite its significant role and the high prevalence of polyploids in nature, neopolyploids face challenges in establishing themselves (Fowler & Levin, 1984; Köhler et al., 2010). Neopolyploids are subjected to strong positive frequency-dependent selection, termed “minority cytotype exclusion”, as they compete with locally adapted parental diploids (Levin, 1975). Overcoming this handicap, which determines the establishment and persistence of neopolyploids, requires mechanisms such as exploiting competitive advantages or occupying unfilled ecological niches (Hijmans et al., 2007; López-Jurado et al., 2019; Ståhlberg, 2009). In addition, previous studies have shown that newly formed polyploid species experience higher extinction rates than their diploid counterparts (Mayrose et al., 2011). This indicates that even after short-term establishment, only a few polyploids persist in the long term (Arrigo & Barker, 2012) normally due to their small population sizes and restricted distributions (Anneberg et al., 2023; Van De Peer et al., 2017). In addition, another possible explanation for this low long-term success of polyploids is the interaction with their ancestral cytotype in zones of emerging secondary contact (Mortier et al., 2024), where previously separated, differentiated polyploids are brought into contact with diploid cytotypes (Mráz et al., 2012; Singliarova et al., 2019). Once in contact, the coexistence of polyploids and diploids is influenced by their different adaptive potentials and competitive interactions (Kolář et al., 2017). In this context, modern coexistence theory (MCT), a framework used for predicting plant species coexistence (Chesson, 2000), have been also applied to examine competition across different environmental conditions (Spaak et al., 2021). MCT can provide insights into the establishment process in the early stages of polyploid evolution (Castro et al., 2023). Additionally, MCT can help estimate the long-term fate of established polyploid species and their competitive interactions with diploid parents (Rey et al., 2017). MCT predicts the coexistence of two cytotypes based on two mechanisms: stabilizing mechanisms, which increase niche difference, and equalizing mechanisms, which reduce the average fitness gap between cytotypes (Adler et al., 2007; Chesson, 2000). In the context of intercytotypic competition, niche differences would represent the average extent to which cytotypes restrict individuals of their own cytotype compared to those of other ploidy levels. Moreover, fitness differences underlie competitive dominance and, in the absence of niche distinctions, determine competitive superiority between pairs of cytotypes.

The competitive dynamics between cytotypes can be influenced by various factors. Niche and fitness differences are not static, as they depend on the abiotic conditions under which cytotypes compete (Germain et al., 2018). Consequently, competitive dynamics can change with shifting climate conditions (Van Dyke et al., 2022). Indeed, climate has been recognized as one of the most influential predictors of the distribution of polyploid species globally (Rice et al., 2019). Thus, understanding the impact of the environment on cytotype coexistence mechanisms is particularly significant under global change scenarios, where climatic conditions are projected to undergo drastic alterations. This is especially relevant in the Mediterranean region, where an increase in temperature is predicted to be accompanied by severe and more frequent drought episodes over the coming years (Giorgi & Lionello, 2008).

Another factor influencing competitive interactions, alongside the environment, is the divergence of functional traits between cytotypes. Polyploidization often induces significant changes in cell size and adjustments in gene regulatory networks (Ebadi et al., 2023), leading to morphological and physiological alterations with substantial ecological and evolutionary implications (Li et al., 1996; Weiss-Schneeweiss et al., 2013). Genome duplication results in larger stomatal guard cells and lower stomatal density, which would positively impact photosynthetic rates (Maherali et al., 2009; Vyas et al., 2007; Zhang et al., 2020). Moreover, polyploids tend to have a lower osmotic potential and a thicker epidermis (Correia et al., 2023; Li et al., 1996), which generally increases drought resistance, potentially conferring polyploids a competitive advantage over diploids in water-limited scenarios (Aschehoug et al., 2016; Kraft et al., 2015). This variation in functional traits may influence fitness so that trait divergence is expected to underlie the nature and magnitude of competitive interactions between polyploids and diploids. Previous studies have investigated the relationship between functional traits and competitive interactions (Cadotte, 2017; Kraft et al., 2015; Kunstler et al., 2016; Pérez-Ramos et al., 2019). Indeed, the specific leaf area (SLA), an effective functional trait widely used as a proxy for plant growth strategy (Quétier et al., 2007; Wright et al., 2004), is globally correlated with low competitive effect (Kunstler et al., 2016). However, the contribution of functional traits to competitive dynamics in the context of polyploidy has been understudied (but see Guo et al., 2023 and Rey et al., 2017). Therefore, it would be valuable to elucidate whether divergence in functional traits or the traits themselves modify competitive interactions between cytotypes under contrasting water availability scenarios.

The autopolyploid *Dianthus broteri* complex (Caryophyllaceae) is an excellent system to unravel the competitive dynamics between polyploids and diploids and the modulating impact of water availability and functional traits. Endemic to the Iberian Peninsula, this polyploid complex comprises four cytotypes (2x, 4x, 6x, and 12x) that form independent monophyletic lineages and are distributed in monocytotypic populations with disjunct geographic ranges (Balao et al., 2009, Balao et al., 2011). Their distinct ecological niches are distributed along an aridity gradient, with the highest ploidy levels (6x and 12x) occupying the most restricted and extreme habitats (López-Jurado et al., 2019). The presence of monocytotypic populations in *D. broteri*, along with a single known secondary 4x-12x contact zone (Balao et al., 2009), indicates an ongoing unstable coexistence between cytotypes. Moreover, cytotypes exhibit incipient divergence in vegetative and reproductive traits (Balao et al., 2011), photochemical responses (López-Jurado et al., 2020), temperature tolerances (López-Jurado et al., 2024) and ecological strategies (López-Jurado et al., 2022). Specifically, 6x and 12x showed the greatest divergence in terms of ecological strategies where 6x showed a nutrient and water conservation strategy (i.e., low SLA) and 12x showed a nutrient and water acquisition strategy (i.e., high SLA). Therefore, examining the way these and other physiological functional traits influence competitive dynamics between cytotypes could provide fundamental insights into the mechanisms underlying these interactions, especially in the context of regional water deficits.

The present study examines the coexistence of distinct cytotypes within common environments, focusing on the impact of water availability and trait divergence. We hypothesize that water availability would modify the competitive dynamics among cytotypes. Specifically, cytotypes with lower ploidy levels (2x and 4x) would dominate under HWA, while cytotypes with higher ploidy levels (12x and 6x) would dominate under LWA. Consequently, the coexistence of cytotypes will be affected too. Furthermore, both trait divergence and inherent traits will influence the competitive interactions among cytotypes. To test these hypotheses, we (1) describe changes in the competitive ability of cytotypes under two contrasting soil water availability scenarios, (2) predict coexistence between cytotypes by analyzing fitness and niche differences, and (3) elucidate potential linkages between functional traits and competitive interactions, thereby providing insights into their competitive interactions under arid conditions.

## Material & Methods

### Experimental competition set-up

We sowed 345 seeds, randomly pooled from three populations of each *D. broteri* cytotype, to ensure representativeness and reduce maternal origin effects. The seeds were germinated in Petri dishes in a chamber model F-1 (Ibercex, Spain), under a 12/12 h photoperiod and 25/17 °C. Once germinated, the seedlings are arranged according to ‘minimal target-neighbour’ design (Hart et al., 2018) to quantify cytotype competition by pairing two individuals in each combination. Thus, we used 14 competitive combinations: 2x-2x, 2x-4x, 2x-6x, 2x-12x, 4x-4x, 4x-6x, 4x-12x, 6x-6x, 6x-12x, 12x-12x including four monospecific stands 2x, 4x, 6x, and 12x (growing alone) under two contrasting water availability regimens. Under high water availability (HWA), soil moisture was maintained between 40% and 45% of the soil water content (% VWC), while under low water availability (LWA), soil moisture was maintained between 15% and 20% VWC. This setup allowed us to examine the effects of intraploidy and interploidy competition within the *D. broteri* complex, and how these ploidy interactions could vary by the availability of soil water.

We assigned 240 seedlings from each cytotype to the 28 experimental combinations (14 competitive combinations × two water availability treatments) and grown in 650 mL pots filled with a mixture of a commercial organic substrate (Gramoflor GmbH und Co. KG.) and perlite (3:1) and placed in a greenhouse with controlled minimum and maximum temperature of 16-26 °C, 40-60 % relative humidity and unfiltered natural light conditions (approximately 14/10 h day/night and maximum photosynthetic photon flux density incident on leaves of 1000 μmol m^−2^ s^−1^). During the experiment, VWC of each pot was measured regularly using an ML3 ThetaProbe soil moisture sensor (Delta-T Devices Ltd) to check soil water levels. In the first two weeks of the experiment, 154 seedlings died due to non-competitive causes, and the competition replicates were excluded, resulting in a final count of 70-86 seedlings from each cytotype.

### Functional trait measurements

After six months of growth under the previously described conditions, six replicates from each competitive combination survived, totalling 168 competitive units. The total dry biomass (above- and belowground) of each individual (n = 277) was quantified after drying for 48 h in a fan-forced oven at 65°C. Additionally, we measured a set of functional traits from each competition combination (apart from the monospecific stands). We performed instantaneous gas exchange measurement in a randomly selected, fully developed leaf per individual (n = 242) using an IRGA gas exchange meter (LI-6800, LI-COR Inc., Neb., USA). We quantified the net photosynthetic rate (A_N_), stomatal conductance (g_s_) and intrinsic water use efficiency (_i_WUE) under the follow leaf cuvette conditions: 400 μmol CO_2_ mol^−1^, 25/28 °C, 50 ± 5% relative humidity, and a photon flux density of 1000 μmol photon m^−2^ s^−1^ (with 15% blue light to maximize stomatal aperture). Furthermore, we investigated the maximum quantum efficiency of PSII photochemistry (F_v_/F_m_) using a portable modulated fluorometer (FMS-2, Hansatech) with dark-adapted leaves (n = 263) measured at noon (1600 μmol photon m ^−2^ s ^−1^). In addition, specific leaf area (SLA; leaf area per unit leaf dry mass) and leaf dry matter content (LDMC; leaf dry mass per unit leaf fresh mass) were determined (n = 380) following the protocols described by Cornelissen (2003). Leaf area was measured using ImageJ software (US National Institutes of Health, Bethesda, MD, USA) from scanned leaves and samples were oven-dried at 65 °C for 48 hours.

### Competitive interactions and the effect of water availability on the MCT

We estimated different competition parameters in the MCT framework (Chesson, 2012) using the *cxr* package ver. 1.0.0 (García-Callejas et al., 2020) in R ver. 4.3.3 (R Core Team, 2024). Our experiment was designed to parameterize a mathematical model describing the population dynamics of interacting species (Ricker, 1954), which was extended here to cytotypes. We employed a minimal target-neighbour experimental design (pairing a focal individual with a competitor). Although a range of competitor densities is typically recommended to improve model accuracy (Hart et al., 2018), simulation studies confirmed the validity of our approach (see Supplementary Methods; Appendix). This population dynamic model, based in the per-capita population growth rate, incorporated the competitive interactions between cytotypes (α_ij_ and α_ii_ for inter- and intracytotypic competition respectively) and their demographic parameters (Godoy & Levine, 2014; Levine & HilleRisLambers, 2009). Detailed mathematical explanations for these parameters are provided in Supplementary Methods. The per capita population growth rate is ideally quantified on an annual basis using recruitment numbers. However, due to the perennial nature of *D. broteri* and the controlled greenhouse conditions of this study, total dry biomass over six months was used as a fitness proxy. Vegetative biomass is positively correlated with fecundity and fitness in many plant species (Younginger et al., 2017), and is widely accepted as a fitness surrogate (Dostál, 2023; Zhang & van Kleunen, 2019). In the related *Dianthus sylvestris*, plant size predicts flowering probability and survival (Pålsson et al., 2024), while in 12x *D. broteri*, vegetative biomass over eight months is strongly correlated with reproductive biomass (R² = 0.85, p-value < 0.001, n = 9).

Based on this model, we estimated the intrinsic growth rates, stabilizing niche differences, average fitness differences, and predicted competitive outcomes. Negative niche differences are indicative of priority effects, which implies that cytotypes are experiencing positive density dependence and suggesting that the first species to arrive in a community gains an advantage. Niche differences between 0 and 1 reflect the degree to which cytotypes limit each other compared to themselves. Furthermore, when niche overlap is complete, the dominant cytotype is determined solely by its average fitness ratio. Fitness differences were derived from two components: the demographic ratio and the competitive response, which represent alternative routes for achieving competitive dominance (Godoy & Levine, 2014). The demographic ratio refers to the relative intrinsic growth rates between cytotypes, while the competitive response refers to the sensitivity of a cytotype to both intracytotypic and intercytotypic competition. We tested the effect of water availability on niche and fitness differences for each cytotype combination using two-tailed paired Student’s t-tests. Three possible outcomes for cytotype coexistence were considered based on these differences: stable coexistence (when niche differences exceed fitness differences), competitive exclusion (when fitness differences exceed niche differences), and priority effects (when niche differences are negative and it is predicted that the first cytotype to arrive in a community gains an advantage(Ke & Letten, 2018). Lastly, from the fitness difference parameter, we derived the competitive ability (Hart et al., 2018) for each cytotype, which reflects the fitness of each cytotype against all other competitors assuming null niche differences (see Supplementary Methods for details). A higher competitive ability can arise from the ability to achieve a large biomass in the absence of competition (i.e., higher intrinsic growth rate) and/or from a reduced sensitivity to competition with other cytotypes (i.e., the geometric mean of α_ij_ and α_ii_).

### Divergence of functional traits in polyploids and its role in competition

To evaluate the divergence in functional traits between cytotypes and the effects of competition (presence/absence) and water availability (HWA/LWA) on these traits, we used generalized linear models (GLMs) with a Gaussian error structure and an identity link function for each dependent variable. Main effects and interactions of competition and water availability were analyzed using Type III sums of squares. Models were fitted to examine the independent effects of each factor and their two-way interactions. Mean trait values for significant factors were compared with Tukey’s post hoc tests (α = 0.05) using the *lsmeans* package in R (Lenth, 2016). Additionally, we explored the complex relationships between traits and competition by expanding the Ricker competition model. We adapted the neighbourhood modelling framework from Kunstler et al. (2016) to our greenhouse experimental setup. In our model, the growth of the focal cytotype (f) is represented as the product of its maximum potential growth (which includes the effects of traits on growth), and the reductions caused by competition from the neighbouring cytotype (c). To quantify this phenomenon, we used the following equation:

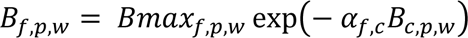

Here, *B*_*f,p,w*_ represents the biomass of the focal individual (f) in competition while *Bmax*_*f,p,w*_is its maximum biomass in the absence of competitors modelled with a normally distributed random effect of ploidy level (*εp*), in which *εp* ∼ N(0, *σ_γ_*), and random effect of watering (*εw*), in which *εw* ∼ N(0, *σ_γ_*). The competitive parameter, *α_f,c,_* is the effect of the neighbour cytotype (c) on the growth of the focal cytotype (*f*), which is multiplied by the biomass of the competitor (*B*_*c,p,w*_). We modelled the effect of traits on the growth model parameters individually, assessing one trait at a time. The effect of a focal cytotype trait value (*t_f_*) on its maximum growth was represented as *m_0_ + m_1_ t_f_*. Here, *m_0_* denotes the average maximum growth, and *m_1_* represents the effect of the focal cytotype trait. The competitive parameter (*α_f,c_*) was modelled using an equation of the form:

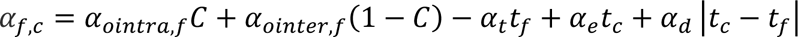

In which *α_ointra,f_* and *α_ointer,f_* are (respectively) trait-independent intraspecific and trait-independent interspecific average competition for the focal cytotypes (*f*), each modelled with a normally distributed random cytotypes effect (*ε*_*f*_) as: *α_ointra,f_* = *α_ointra,f_* + *εf*. The parameter *C* is a binary variable that takes the value 0 for intercytotypic combinations and 1 for intracytotypic combinations; *α*_*t*_ represents the competition tolerance (or competitive response) of the focal species, indicating the alteration in competition tolerance attributed to the traits (*t*_*f*_) of the focal individual; *α*_*e*_ represents the competitive effect, denoting the alteration in competition effect caused by the traits (*t*_*c*_) of the neighbouring individual, while *α_d_* signifies the effect of trait dissimilarity, indicating the change in competition due to the absolute distance between traits |*t*_*c*_ − *t*_*f*_|. Finally, it should be noted that values of *α_f,c_* > 0 indicate competition, while *α_f,c_* < 0 indicates facilitation.

To estimate the standardized coefficients for each model, both the response and explanatory variables were standardized by dividing them by their respective standard deviations before analysis. We conducted two versions of each model. In the first version, the parameters (m_0_, m_1_, α_0_, α_t_, α_e_, α_d_) were estimated as constants across water availability treatments (HWA and LWA). In the second version, we allowed for different fixed estimates of these parameters for each water availability, enabling us to examine the variation between the two experimental conditions more closely (Kunstler et al., 2016). The trait-based neighbourhood models were fitted using dynamic Hamiltonian Monte-Carlo sampling in Stan v.2.32.2 (Stan Development Team 2024), with the package *rstan* v.2.32.6 as a front-end. We fitted the growth model running five chains of 10,000 iterations, including 5,000 iterations of adaptation. Weakly informative priors were used for all parameters. The main parameters m_0_, m_1_, *α*_*intra*_ and *α*_*inter*_ for the HWA and LWA conditions were assigned priors with a mean of 0 and a standard deviation of 1 (see Stan code for the rest of priors).

## Results

### Competitive interactions are largely affected by water availability

Our results showed that soil water availability influenced the competitive abilities of cytotypes (Fig. 1 a, b). Under HWA (Fig. 1a), the tetraploid cytotype (4x) exhibited the highest competitiveness, followed by the hexaploid (6x), diploid (2x), and dodecaploid (12x). However, under LWA, this order was reversed. The 6x cytotype demonstrated the highest dominance, followed by the 12x, 2x, and 4x, respectively (Fig. 1b).

**Fig. 1:**
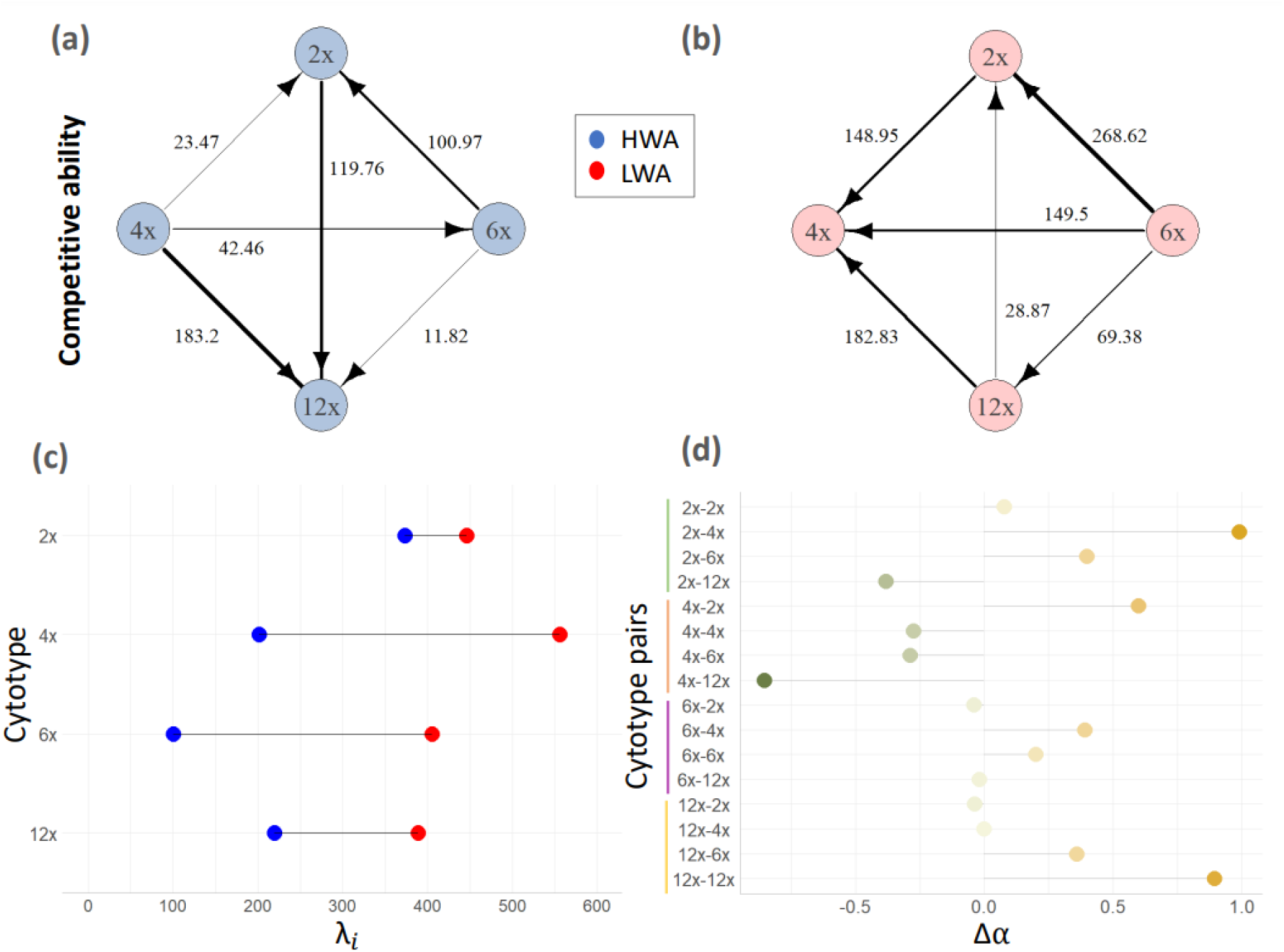
Competitive ability of *Dianthus broteri* cytotypes (2x, 4x,6x and 12x) under both high **(a)** and low water availability **(b)** conditions. The direction of the arrow indicates the cytotype with the lowest competitive ability in each pair (i.e. the worst competitor within each pair), while the thickness of the arrow denotes the magnitude of fitness differences. Changes in the direction of the arrows indicate changes in the dominance between the two irrigation scenarios. **(c)** Intrinsic growth rate (λ_i_) of the different cytotypes in the absence of competition (blue dots indicate HWA conditions, while red dots indicate LWA conditions). **(d)** Difference in interaction coefficients between HWA and LWA conditions (Δα = α_HWA_ – α_LWA_). Positive differences (represented by orange dots) indicate a greater interaction coefficient for that pair under HWA compared to LWA, whereas negative differences (green dots) indicate the reverse pattern.

When competitive ability was decomposed into intrinsic growth rate (i.e., biomass in the absence of competition) and sensitivity to competition (i.e., the geometric mean of intra- and interspecific interaction coefficients), we observed a significant increase in intrinsic growth rates across all cytotypes under LWA (paired t-test t = 3.5137, df = 3, p-value <0.05 Fig. 1c). However, the dominant cytotype (with the highest competitive ability) under each water availability condition (4x under HWA and 6x under LWA) did not exhibit the highest intrinsic growth rate compared to other cytotypes, but rather showed a lower sensitivity to competition. The water availability markedly altered the intensity of both intra-(α_ii_) and interspecific (α_ij_) competition but those differed between cytotype pairs (Fig. 1d). The tetraploids under HWA had overall remarkably low α_ii_ and α_ij_ (except for their interaction with diploids), which explains their high competitive ability. Conversely, the tetraploids exhibited the lowest competitive ability under LWA as a consequence of increased interaction coefficients (specifically, 4x-12x pair). In turn, the hexaploids showed the highest competitive ability under LWA due to their higher tolerance to competition, as evidenced by their notably low interaction coefficients (Fig. 1d). Additionally, the dodecaploid cytotype showed a substantial disparity in its intraspecific interaction coefficient which was notably higher under HWA compared to LWA. Consequently, the 12x cytotype displayed the lowest competitive ability under HWA due to its increased sensitivity to competition. Moreover, for the diploids, its competitive ability was not affected by the water availability treatment, since in both cases it was intermediate (Fig. 1a).

Similarly, niche difference was significatively higher in HWA than LWA (paired t-test, t = −3.9509, df = 5, p-value <0.05; Fig. 2), whereas the fitness difference showed the opposite pattern (paired t-test, t = −3.5499, df = 5, p-value <0.05; Fig. 2). Nevertheless, the feasibility of cytotype coexistence was rarely predicted with only one pair of cytotypes coexisting in each water availability condition (Fig. 2). Under HWA, our model predicted coexistence of the 4x-12x pair mainly due to high niche differences. In the other combinations (4x-2x and 4x-6x), the tetraploids tended to dominate, excluding other cytotypes. In contrast, under LWA conditions, the model predicted the coexistence of 2x and 4x cytotypes (given their high niche difference and low fitness difference). Meanwhile, the hexaploids outcompeted the other cytotypes –including tetraploids– under LWA, leading to their competitive exclusion. Additionally, the 12x-2x pair exhibited strong priority effects due to negative values in the niche difference (Fig. 2).

**Fig. 2:**
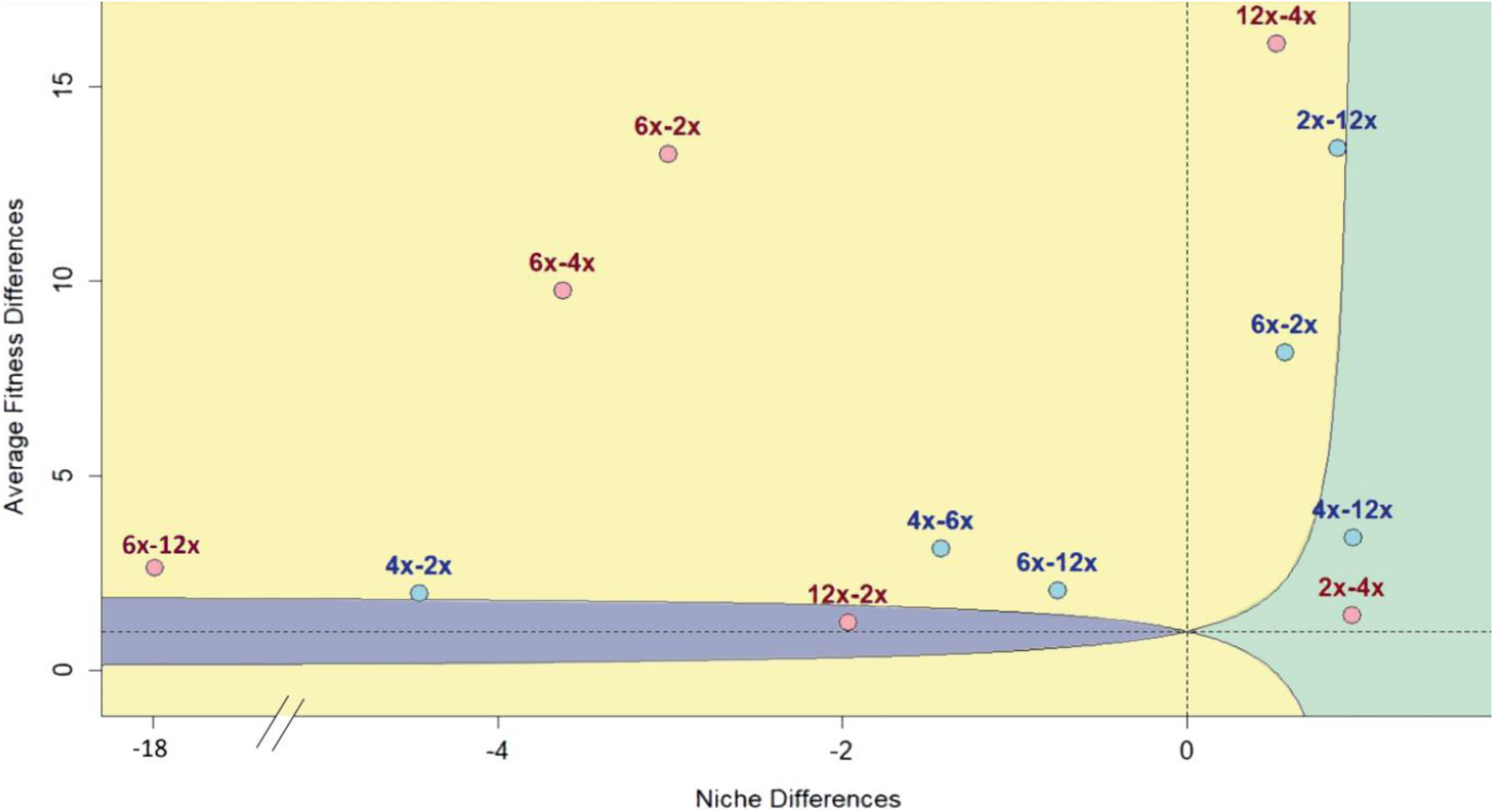
Relationship between fitness and niche differences for each competitive interaction between *Dianthus broteri* cytotypes (2x, 4x, 6x, and 12x) under HWA (blue dots) and LWA conditions (red dots). The three different competitive outcomes are represented as follows: coexistence (green area), where both cytotypes can persist together; priority effects (purple area), where the first cytotype to arrive dominates the competition; and competitive exclusion (yellow area), where the first cytotype excludes the second for a specific combination.

### Functional trait divergence

Our results showed that water availability had a significant impact on all functional traits (Table 1). Specifically, the LWA treatment increased A_N,_ g_s_, LDMC and _i_WUE by 9.13-109.15%. In contrast, SLA and F_v_/F_m_ values were reduced by 8.13-67.8%. Furthermore, cytotypes showed substantial variations in g_s_, SLA, and LDMC, with significant differences being specific to each cytotype pair. We observed significant differences in g_s_ between 4x (112.69 ± 10.23 mmol H_2_O m^−2^ s^−1^) and 12x (154.45 ± 10.49 mmol H_2_O m^−2^ s^−1^; Tukey’s post hoc test; p-value < 0.05). SLA differed between 2x (11.44 ± 0.45 m^2^ Kg^−1^) and 12x (19.96 ± 2.49 m^2^ Kg^−1^; Tukey’s post hoc test; p-value < 0.05). In addition, LDMC showed differences between 2x (309.59 ± 6.78 g) and 6x (259.19 ± 11.04 g; Tukey’s post hoc test; p-value < 0.05) and between 2x (309.59 ± 6.78 g) and 12x (263.60 ± 9.19 g; Tukey’s post hoc test; p-value < 0.05). Alternatively, the interaction between ploidy level and water availability was not observed to be significant for any of the traits. Similarly, competition did not independently affect any functional trait, nor did its interaction with the water availability treatment (Table 1). However, the interaction between competition and ploidy level led to significant divergence in g_s_, with values decreasing for 2x plants in competition (111.08 ± 10.80 mmol H_2_0 m^−2^ s^−1^) compared to when they were grown alone (163.47 ± mmol H_2_0 m^−2^ s^−1^; Tukey’s post hoc test; p-value < 0.05).

**Table 1:** Results of Generalized Linear Models (GLM) analyzing the effect of ploidy level (2, 4x, 6x and 12x), competition (alone or with competitor), water availability (high or low), along with their pairwise interactions, on *Dianthus broteri* functional traits. Degrees of freedom (df) and F-value are shown, with significant results (p-value < 0.05) indicated by an asterisk.

|  | df | A<sub>N</sub> | g<sub>s</sub> | SLA | LDMC | F<sub>v</sub>/F<sub>m</sub> | iWUE |
| --- | --- | --- | --- | --- | --- | --- | --- |
| **Competition** | 1 | 0.2635 | 0.3854 | 0.0426 | 0.0978 | 0.0051 | 0.0222 |
| **Water availability** | 1 | 25.1462* | 25.375* | 14.2780* | 4.9718* | 13.7343* | 9.7650* |
| **Ploidy** | 3 | 1.9683 | 2.8713* | 4.1760* | 6.41* | 1.2733 | 0.5628 |
| **Competition:Water availability** | 1 | 0.0153 | 0.119 | 0.2119 | 0.0791 | 0.6581 | 0.8269 |
| **Water availability:Ploidy** | 3 | 2.5397 | 2.5918 | 2.9411 | 0.4390 | 0.1053 | 0.4435 |
| **Competition:Ploidy** | 3 | 0.6563 | 3.221* | 4.9168 | 2.364 | 0.5432 | 1.1277 |

### Functional traits promote tolerance and competitive effects

Our results indicate that functional traits play a crucial role in shaping competitive dynamics within the *D. broteri* complex, with soil water availability significantly influencing these relationships. The effects of these traits are driven by four main mechanisms. Firstly, LMDC and SLA traits were directly associated with higher and lower growth rates, respectively, in the absence of competition (m_1_, Fig. 3). These global effects were influenced by water availability, being pronounced for SLA under HWA and for LMDC under LWA. Furthermore, lower values of A_N_, F_v_/F_m_ and _i_WUE were associated with increased tolerance to competition from neighbouring cytotypes (i.e., ‘competitive response’; *α_t_t_f_* < 0, Fig. 3). The competitive response of these traits persisted under LWA but not under HWA. Additionally, a stronger competitive effect (α_e_t_c_) would enable cytotypes with higher trait values to suppress the growth of neighbouring cytotypes more effectively, SLA under LWA conditions exerted a significant positive competitive effect (α_e_t_c_ > 0, Fig. 3). Lastly, our analysis revealed a global increased trait dissimilarity effect (α_d_ |t_c_-t_f_|) for A_N_ and LDMC, which led to decreased competition and reduced suppression of focal cytotype growth. This effect was maintained under both water availability conditions for LDMC and under LWA for A_N_ (α_d |tc-tf |_ > 0, Fig. 3).

**Fig. 3:**
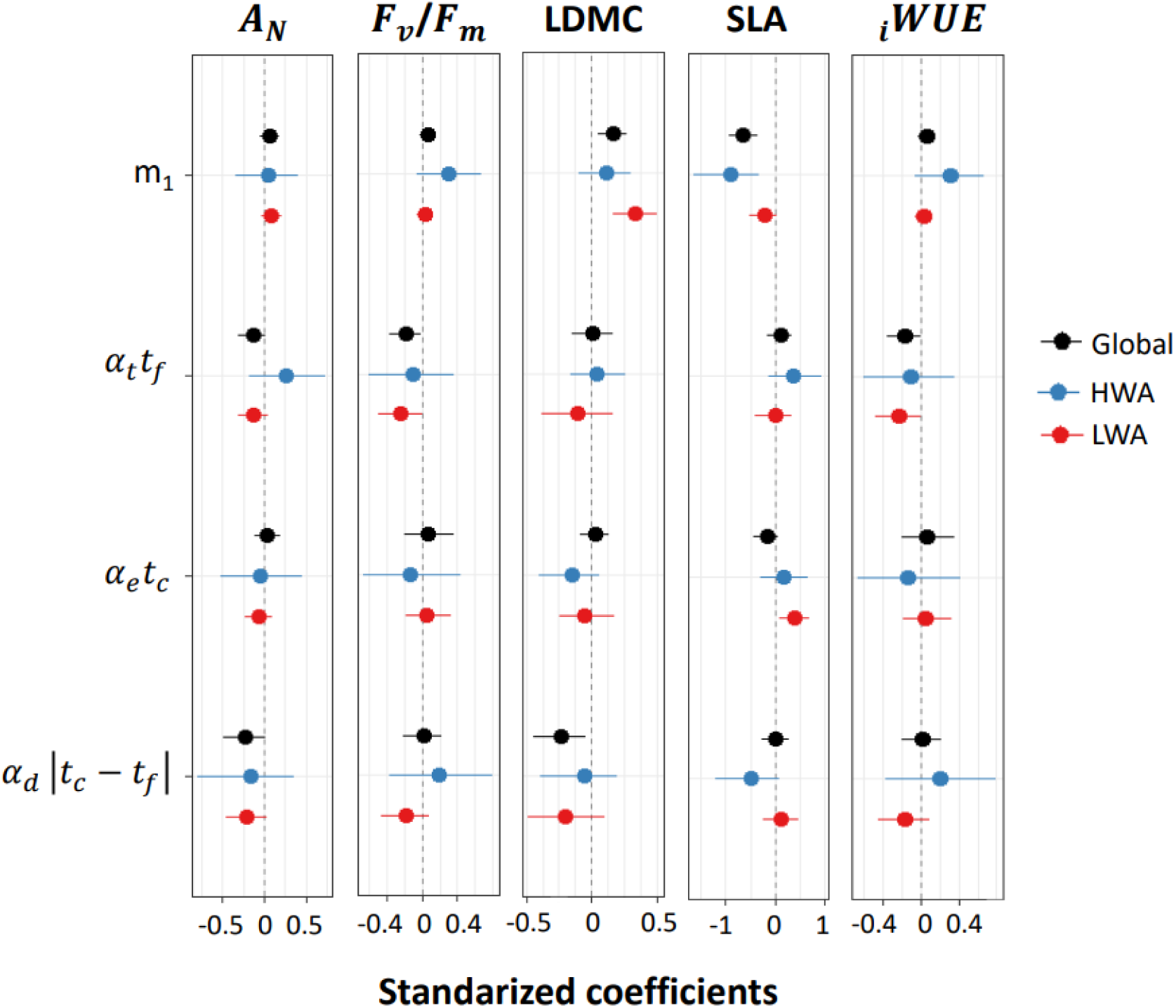
Standardized regression coefficients for growth models, adjusted separately for net photosynthetic rate, *A*_N_, maximum quantum efficiency of PSII photochemistry F_v_/F_m_, leaf dry matter content, LDMC, specific leaf area, SLA and intrinsic water use efficiency _i_WUE. Stomatal conductance (*g*_s_) showed posterior estimates that included zero for all competition parameters. Dots represent average estimates, while lines indicate 95% confidence intervals. Dark dots and lines depict joint control and drought estimates, blue represents HWA conditions, and red represents LWA. Parameter estimates include: the effect of the focal individual trait (*t_f_*) value on maximum growth (m_1_), the effect of the focal cytotype trait value (t_f_) on its tolerance to competition (*α_t_tf_*) (positive values indicate higher trait values result in greater tolerance to competition), the effect of trait values in the individual competitor (t_c_) on its competitive effect (α_e_tc_) (positive values indicate higher trait values lead to greater reduction in focal cytotype growth) and the effect of trait dissimilarity between the focal cytotype and its competitors (α__d |t_c-t_f |_) (negative values indicate greater dissimilarity leads to less reduction in focal cytotype growth). Significant estimates are those that do not cross the zero line.

## Discussion

### Competitive interactions are largely affected by soil water availability

While numerous studies have reported differences in competitive ability between polyploids and diploids, usually based on indirect evidence (Castro et al., 2023; Čertner et al., 2019; Grewell et al., 2016), other studies have found no significant support for these differences (Hülber et al., 2011; Münzbergová, 2007). Our work examines the fine-scale competitive dynamics among ploidy levels in the *Dianthus broteri* complex, focusing on the implications for the long-term persistence of polyploids and their response to variations in soil water availability. We employed recent advancements in ecological coexistence theory and parameterized models, demonstrating that both polyploidy and water availability significantly affected most competition parameters. In line with polyploids generally experiencing delayed ontogenesis and exhibiting a reduced rate of biomass accumulation (Corneillie et al., 2019; Deng et al., 2012), we found here that the intrinsic growth in higher cytotypes (6x and 12x) was consistently lower than in lower ploidy level cytotypes (2x and 4x; Fig 1c). This phenomenon, referred to as “high-ploidy syndrome” (Tsukaya, 2008), has been associated with an elevated demand for resources in lineages after genome duplication (Anneberg & Segraves, 2023; Guignard et al., 2017). However, there are studies that suggest the opposite, indicating that polyploids exhibit a faster growth rate and larger organ size (Šmarda et al., 2023; Sugiyama, 2005).

Additionally, consistent with the known effect of drought on interspecies competition (Adler et al., 2012; López et al., 2013), our results suggest that soil water availability induces changes in the network of competitive interactions within the *D. broteri* complex (Fig. 1a, b). Under HWA, presumably the most benign condition, the tetraploids exhibited the highest competitive ability. This finding aligns with the observation that 4x is the most geographically widespread cytotype (Balao et al., 2009; López-Jurado et al., 2019). Conversely, the highest ploidy levels (12x and 6x) demonstrated the lowest competitive ability, consistent with their restricted distribution to the most arid areas. Congruently, under LWA, the competitive hierarchy was reversed, with 6x and 12x emerging as superior competitors. This finding is supported by previous studies highlighting the adaptation of those higher ploidy *D. broteri* cytotypes to stressful conditions (López-Jurado et al., 2020, 2024) and the role of specific ecophysiological traits (i.e, F_v_/F_m_, relative water content or leaf water potential) in drought tolerance (López-Jurado et al., 2016). While these results are consistent with previous studies indicating that water deficit can enhance the competitive ability of polyploid species over diploids (Guo et al., 2023; Rey et al., 2017), no differences in competitive abilities between diploids and tetraploids were observed under reduced soil moisture in *Chamerion angustifolium* (Thompson et al., 2015).

### Competitive exclusion is mediated by soil water availability

As indicated in the competitive outcomes analysis, our competition model predicted a general pattern of competitive exclusion among *D. broteri* cytotypes under secondary contact. This aligns with the allopatric distribution of the cytotypes (Balao et al., 2009) and their divergence in ecological niches (López-Jurado et al., 2019), suggesting that intercytotypic competition is shaping their current distributions. Similarly, the Mediterranean *Brachypodium distachyon* and other polyploid complexes have exhibited this intercytotype exclusion pattern (Buggs & Pannell, 2007; Ćertner et al., 2017; Rey et al., 2017), while stable coexistence, at least in the short term, has been found in other mixed-ploidy species (Kao & Parker, 2010; Keeler, 2004). In *D. broteri*, soil water availability affected competitive interactions between cytotypes and, ultimately, their predicted coexistence outcomes through modifications in niche and fitness differences, as observed in other studies (HilleRisLambers et al., 2012; Matías et al., 2018; Wainwright et al., 2019). Specifically, under LWA, fitness differences increased, and niche differences decreased, collectively reducing the likelihood of coexistence. Notably, our model predicted the stable coexistence of 4x and 12x cytotypes in HWA, which represents the only secondary contact zone of the complex found in nature (Balao et al., 2009). This stable coexistence would be possible given the poor competitive ability displayed by the rare 12x, due to its strong self-limitation (Fig. 1d; Chesson, 2000; MacArthur, 1970; Mcpeek, 2012). However, under LWA, the 2x-4x coexistence was also predicted and the possibility of current or new stable secondary contact zones should not be dismissed based on their close distributions and overlapping ecological niches (Balao et al., 2009).

Additionally, the role of chance in the long-term of survival of polyploids has been overlooked (Mortier et al 2024). Founders that establish successfully can secure available resources and prevent the establishment of later-arriving cytotypes with similar niche preferences, regardless of their comparative fitness (Kardol et al., 2013). This priority effect ensures the continued dominance of the initially established cytotype, as predicted for the 2x-12x interaction. Nevertheless, these priority effects were again modulated by water availability, hindering the long-term fate predictions. As water availability increases, the risk of local extinction for higher *D. broteri* polyploids (6x-12x) also rises. Conversely, under the predicted future aridity scenario for the Iberian Peninsula, these higher polyploids (6x and 12x) are expected to expand and outcompete lower ploidy cytotypes, ultimately stabilizing in the long term (Ræbild et al., 2024). This scenario is consistent with the non-random establishment and long-term survival of many polyploid lineages, which have coincided with major periods of global climate change and environmental stress (Cai et al., 2019; Van De Peer et al., 2017; Van de Peer et al., 2021).

### Functional traits modulate competitive effects and response

Phenotypic changes resulting from polyploidization, particularly those related to physiological plasticity, can confer fitness advantages and modify competitive effects under water shortage (Guo et al., 2023). Our analysis revealed significant inter-cytotype differences in g_s_, SLA, and LDMC, which is in accordance with previous studies on the *D. broteri* complex (López-Jurado et al., 2022, 2024). Notably, the 4x cytotype exhibited significantly lower g_s_ compared to the 12x, as also observed under heat, suggesting divergent stress response strategies (López-Jurado et al., 2022, 2024). Furthermore, previous studies have suggested that the 12x cytotype employs a physiological avoidance mechanism, evidenced by sufficiently high levels of g_s_ that maintain constant relative water content and leaf water potential (López-Jurado et al., 2016)). This strategy may explain its competitive dominance over the 4x under abiotic stress (Fig. 1ab).

Similarly, our trait-based neighbourhood analyses confirmed the direct and indirect roles of the functional traits mediating competitive interactions (Godoy & Levine, 2014; Kraft et al., 2015). We observed a direct relationship between LDMC and growth rate in the absence of competition under LWA. Higher LDMC values imply thicker and stiffer cell walls. This allows for the maintenance of turgor at lower leaf water potentials (Cheung et al., 2011; Zimmermann, 1978), which may help minimize cell damage during drought (Bowman & Roberts, 1985) and consequently mitigating some of its adverse effects (Blumenthal et al., 2020). Strikingly, despite most studies suggest a strong positive relationship between SLA and the relative growth rate in woody species (Cornelissen et al., 1998; Lambers & Poorter, 1992; Poorter et al., 1990), our results revealed an opposite negative relationship in both HWA and LWA scenarios. This discrepancy may be attributed to the species-specific growth habit variations, as negative relationship between SLA and plant size and height has been observed in herbaceous species (Kleyer et al., 2019). Focusing on our case, we observed that higher-level polyploids (12x and 6x) exhibited a lower intrinsic growth rate compared to lower-level polyploids (2x and 4x; Fig 1d). Additionally, cytotypes showed significant divergence in SLA (Table 1), with higher-level polyploids displaying higher values. This trend may reflect greater investment in leaf surface area by higher ploidy levels, potentially facilitating the release of excess heat through evapotranspiration in more arid habitats (El Nadi, 1974). Therefore, we suggest that polyploidy may be altering patterns of phenotypic covariation (Baker et al., 2024; Balao et al., 2011) by changing the genetic and regulatory bases underlying trait expression, leading to new relationships between traits that were previously predictably correlated. Additionally, functional traits exhibited associations with enhanced competitive effects or improved tolerance to competition, being both predominantly contingent upon water availability. Regarding competitive effects, a modest positive correlation was observed with SLA under LWA, indicating that higher SLA in competitors corresponds to reduced biomass in focal individuals. Species with elevated SLA generally exhibit higher growth rates and shorter lifespans, characteristic of a resource-acquisition strategy on the leaf economics spectrum (Poorter et al., 2009; Wright et al., 2004). This acquisitive strategy, as manifested by competitors, would restrict resource availability for the growth of the focal individual. Furthermore, higher F_v_/F_m_ and A_N_, related to the efficiency and capacity of photosynthesis, were correlated with lower tolerance to competition (*α_t_t_f_*; Fig. 4). This negative relationship suggests a trade-off between the ability of cytotypes to tolerate competition and their photosynthetic performance in the absence of resource competition. Although several studies have documented this trade-off, the underlying mechanisms remain to be elucidated (Campbell & Holdo, 2017; Cope et al., 2021). Similarly, _i_WUE also exhibited a negative relationship with tolerance to competition. Plants with high _i_WUE are typically associated with a resource-conservative strategy, where they prioritize longevity over rapid growth (Bacelar et al., 2012). In contrast, in highly competitive environments with limited water resources, plants that adopt an acquisitive strategy, characterized by lower _i_WUE, often have an advantage because they maximize rapid resource uptake (Wright et al., 2004). As a result, plants following a conservative strategy (high _i_WUE) may exhibit lower tolerance to competition because they are less able to respond dynamically to resource availability in competitive environments.

Lastly, trait dissimilarity had little effect on competition between *D. broteri* cytotypes. Dissimilarity in net photosynthetic rate (A_N_ *α_d_ |t_c_-t_f_|*) led to decreased competition, supporting the “competition-trait similarity” hypothesis, which predicts that the intensity of competition between individuals or species increases with the similarity of their ecological and physiological traits (Cadotte et al., 2013; Falster & Westoby, 2003; Wong et al., 2021). Greater differences in functional traits between species would indicate greater niche differentiation (D’Andrea & Ostling, 2016), allowing species to minimize competition and partition the local resource pool despite overlapping in resource use and ecological preferences (Macarthur & Levins, 1967).

### Concluding remarks

Our findings underscore that the long-term fate of polyploids is highly dependent on environmental variation, which modulates the strength of competitive interactions and coexistence. Water availability in the soil influences ecological settings such as niche and fitness differences, determining competitive outcomes between cytotypes in *D. broteri*. We predicted pairwise-cytotype competitive exclusion, leading to the observed single-ploidy populations and the displacement of higher cytotypes to arid areas. Dominance of higher ploidies under water-deficit conditions is predicted, supporting the polyploid advantage in these stressful environments. Additionally, we elucidate links between functional traits related to leaf physiological performance (i.e., A_N_, F_v_/F_m_, _i_WUE and SLA) and tolerance and competitive effect parameters underlying competitive outcomes. Our findings offer valuable insights into how environmental conditions, ploidy levels and their associated functional traits interact to shape ecological relationships and cytotypes distributions, as well as enhance our understanding and prediction of long-term establishment and persistence of polyploids. Provided that these results can be extended to other polyploid complexes with similar growth habits in the Mediterranean region, our study offers important new mechanistic insights into their competitive dynamics.

## Acknowledgments

We thank to MF, CM, JPA and AT for contributing to the experiment setup. As well as, to the University of Seville Greenhouse and Herbarium General Research Services (CITIUS) for assistance and providing facilities and equipment. Financial support was provided by grants PGC2018-098358BI00 (Spanish Ministerio de Ciencia e Innovación), PDI2023-147752NBI00 and FEDER-US-1381232 (supported by FEDER funds, Andalusian Government and University of Seville), as well as a Margarita Salas post-doctoral fellowship to J. López-Jurado (Spanish Ministerio de Universidades/NextGenerationEU; USE-24438-N) and a FPU pre-doctoral grant to A. Rodríguez-Parra (Spanish Ministerio de Universidades; FPU22/00818).

## Author contributions

FB conceived the study and designed the experiment. ARP and FB performed the statistical analyses and modelling. ARP, FB, JLJ and EMN interpreted the results. ARP and FB wrote the first draft of the manuscript. All authors critically contributed substantial revisions to the manuscript and approved the final version.

## Data availability

The data that support the findings of this study and the datasets and code for the method developed here are available on GitHub https://gitfront.io/r/Albarp13/8CULZbE5v3Ec/Dianthus-broteri-Competition/

