## Supplementary material for "Functional traits and soil water availability shape competitive interactions in a diploid-polyploid complex": Suplementary Methods and Appendix

### Supplementary Methods

Our experiment was designed to parameterize a mathematical model describing the population dynamics of interacting species (Ricker, 1954), which was extended here to cytotypes. This model incorporates the demographic parameters of plants and their competitive interactions, allowing for the quantification of stabilizing niche differences, average fitness differences, and predicted competitive outcomes (Godoy & Levine, 2014;

Levine & HilleRisLambers, 2009). The model is described as follows:

$$\ln(B_i) = \ln(\lambda_i) - \alpha_{ii}\,N_i - \alpha_{ij}N_j - \alpha_{ik}\,N_k \ldots - \alpha_{iz}\,N_z$$

In this context, $B_i$ is the individual biomass of the target cytotype $i$, $\lambda_i$ is the intrinsic growth rate of the target cytotype (biomass of cytotype $i$ in the absence of competition is independent of population density), $\alpha_{ii}$ is the intraspecific competition coefficient describing the per individual effect of the cytotype $i$ on itself, $N_i$ is the number of planted individuals of species $i$ (which is always 1 in our case) and $\alpha_{ij}$ is the interaction coefficients which describe the per capita effect of cytotype $j$ on cytotype $i$.

As pointed out by (Zhang & van Kleunen, 2019), ideally, $B_i$ should be $\frac{N_{i,t+1}}{N_{i,t}}$, the per capita population growth rate from one year to the next based on the number of recruits. However, estimating the population size of perennial plants that reproduce sexually in greenhouses is difficult. Therefore, dry biomass (both above and belowground) was used as an alternative proxy for growth parameter.

Building upon the competition dynamics among cytotypes outlined in this population model, we employed the methodology proposed by Chesson (2012) to assess fitness and niche distinctions between species pairs. Subsequently, following the methodology detailed by (Godoy & Levine, 2014) we calculated firstly the niche overlap (ρ) between cytotype pairs:

$$\rho = \sqrt{\frac{\alpha_{ij}}{\alpha_{jj}} \cdot \frac{\alpha_{ji}}{\alpha_{ii}}}$$

Niche overlap represents the average extent to which cytotypes restrict individuals of other ploidy levels ($\alpha_{ij}$, $\alpha_{ji}$) compared to those of their own cytotype ($\alpha_{ii}$, $\alpha_{jj}$). It ranges from zero (i.e., no overlap) to infinity, indicating that niche differences (expressed as 1 – ρ) can range from negative infinity to one. Negative niche differences are indicative of priority effects, suggesting that the first species to arrive in a community gains an advantage. Niche differences between 0 and 1 reflect the degree to which species A and B limit each other compared to themselves.

Secondly, we proceed to calculate the fitness differences that underlie competitive dominance and, in the absence of niche distinctions, determine competitive superiority between pairs of cytotypes. We define average fitness differences between each pair of competing cytotypes are defined as follows:

$$AFD_i = 1 - \left(\frac{\log(\lambda_i)}{\log(\lambda_j)}\right) \sqrt{\frac{\alpha_{ji}}{\alpha_{ii}} \cdot \frac{\alpha_{jj}}{\alpha_{ij}}}$$

This equation reveals that the fitness difference ($AFD_i$) arises from the interplay of two main factors: the 'demographic ratio' and the 'competitive response'. The first term quantifies the extent to which cytotype *j* outperforms cytotype *i* in biomass production in the absence of competition. The second component elucidates the degree to which cytotype *i* is more susceptible to both intra- and interspecific competition compared to cytotype *j*. Consequently, a species may demonstrate superior competitiveness either by having a large biomass without competition or by being less affected by competition. To assess the fitness of each species against all other competitors, we quantify $AFD_i$ as an absolute quantity, rather than as part of a proportion, assuming zero niche differences. So, starting from the $AFD_i$ expression, we can describe the competitive ability of the cytotype (Hart et al., 2018):

$$K_i = \frac{\lambda_i}{\sqrt{\alpha_{ij}\,\alpha_{ii}}}$$

Competitive ability ($K_i$) refers to a cytotype's capacity for outcompeting others, which can stem from its ability to achieve a large biomass in the absence of competition ($\lambda_i$) or its sensitivity to competition with other cytotypes. According to Chesson (2012), coexistence requires both species to invade when rare and this condition is satisfied when (Godoy & Levine, 2014):

$$\rho < \frac{K_j}{K_i} < \frac{1}{\rho}$$

Based on this condition, we can identify three potential coexistence outcomes. Firstly, stable coexistence arises when niche differences exceed fitness differences. Secondly, competitive exclusion occurs when fitness differences outweigh niche differences. Lastly, priority effects emerge when niche differences are negative, suggesting positive density dependence experienced by species.

**Appendix:**

To evaluate the suitability of our minimal target-neighbour design (one focal individual paired with one competitor) for fitting Ricker's population dynamics model, we conducted 100 simulations incorporating varying competitor densities (1:2:4:8:16) across four species (Sp1, Sp2, Sp3, and Sp4) using the *cxr* package with the same settings applied to our data (see Materials and Methods). We predefined intrinsic growth rate ($\lambda_i$ =219) and interaction coefficients ($\alpha_{11}$ = 0.999, $\alpha_{12}$ = 0.6, $\alpha_{13}$ = 0.01 and $\alpha_{14}$ = 0.876), with ten replicates per competitive combination, to generate fitness values as inputs for the Ricker model. This approach allowed us to compare the estimates of intrinsic growth rate of Sp1 (λ) and interaction coefficients ($\alpha_{11}, \alpha_{12}, \alpha_{13}$ *and* $\alpha_{14}$ ) across three scenarios: the model with one density (the minimal target-neighbour design, d1), the model with three densities (d3), and the model with five densities (d5).

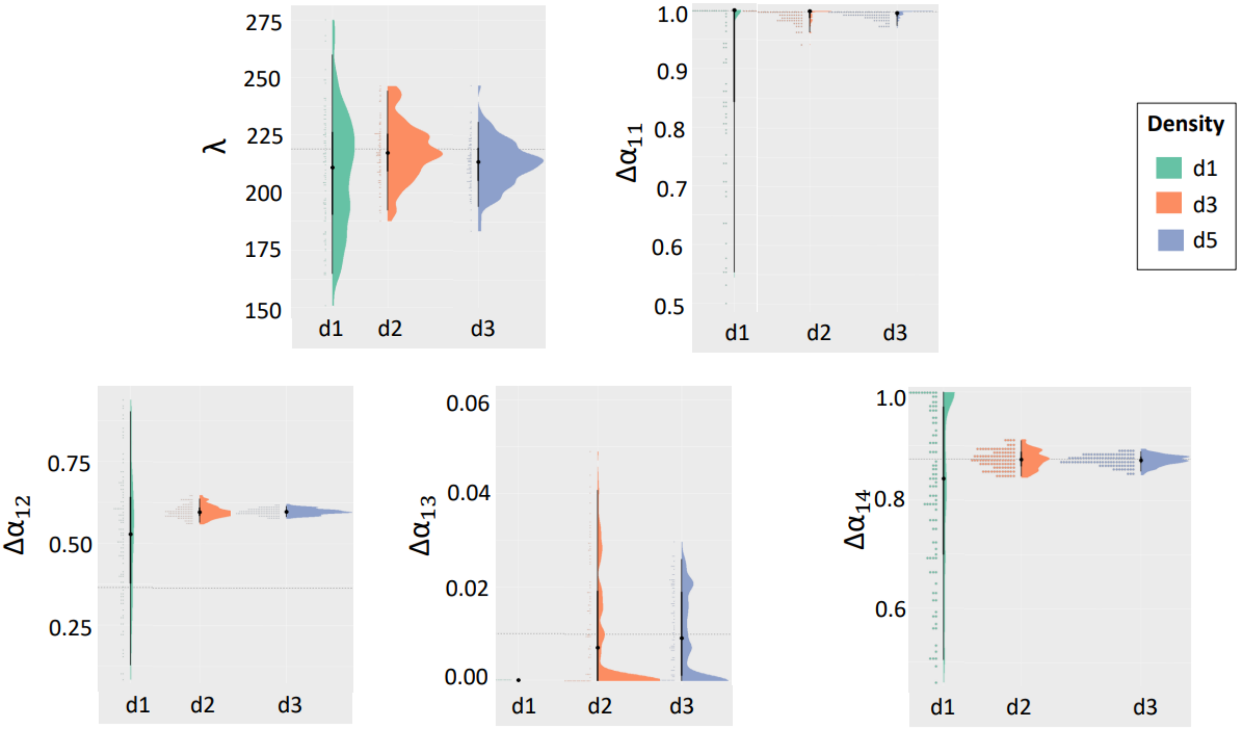
